# BLM helicase suppresses recombination at G-quadruplex motifs in transcribed genes

**DOI:** 10.1101/173252

**Authors:** Niek van Wietmarschen, Sarra Merzouk, Nancy Halsema, Diana C.J. Spierings, Victor Guryev, Peter M. Lansdorp

## Abstract

Bloom syndrome is a cancer predisposition disorder caused by mutations in the *BLM* helicase gene. Cells from persons with Bloom syndrome exhibit striking genomic instability characterized by excessive sister chromatid exchange events (SCEs). We applied single-cell DNA template strand-sequencing (Strand-seq) to map the genomic locations of SCEs at a resolution that is orders of magnitude better than was previously possible. Our results show that, in the absence of BLM, sister chromatid exchanges in human and murine cells do not occur randomly throughout the genome but are strikingly enriched at coding regions, specifically at sites of putative guanine quadruplex (G4) motifs in transcribed genes. We propose that BLM protects against genome instability by suppressing recombination at sites of G4 structures, particularly in transcribed regions of the genome.

## INTRODUCTION

Bloom syndrome (BS) is a rare genetic disorder caused by mutations in the BLM gene, which encodes for the BLM helicase (German, 1993). Symptoms of the disease include short stature, immunodeficiency, UV sensitivity, reduced fertility, and a strong predisposition towards a wide range of cancers. Cells from BS patients display marked genome instability, characterized by elevated spontaneous mutations rates (Vijayalaxmi et al., 1983; Warren et al., 1981), a high frequency of micronuclei (Yankiwski et al., 2000), increased DNA breaks during S-phase (Li et al., 2004), high levels of replication intermediates (Lonn et al., 1990), delayed DNA chain growth during replication (Hand and German, 1975), and an increased sensitivity to DNA damaging agents such as mitomycin C (Hook et al., 1984), ethyl methanesulfonate (Krepinsky et al., 1979), hydroxyurea (Davies et al., 2004), formaldehyde (Kumari et al., 2015), UV radiation (Krepinsky et al., 1980), X-radiation (Kuhn, 1980), and γ-radiation (Aurias et al., 1985). The main cellular phenotype, as well as a diagnostic criterion for Bloom Syndrome, is a marked ~10-fold increase in the rate of sister chromatid exchange events (SCEs) in cells from patients compared to healthy controls (Chaganti et al., 1974; van Wietmarschen and Lansdorp, 2016).

SCEs are a by-product of double strand breaks (DSBs) or collapsed replication forks that are repaired via homologous recombination (HR) (Painter, 1980; Wu, 2007). The HR pathway uses homologous sequences on sister chromatids for repair without loss of genetic information (Li and Heyer, 2008). However, this type of repair requires a physical crossover of DNA strands in the form of a Holliday junction, which needs to be dissolved by BLM, along with its partners TOPO3a, RMI1, and RMI2, to prevent SCE formation (Karow et al., 2000; Wu and Hickson, 2003). BLM also promotes regression of stalled replication forks, thus facilitating fork restart and preventing fork collapse and the formation of DSBs (Davies et al., 2007; Machwe et al., 2006). BS cells display higher numbers of γH2Ax foci (Rao et al., 2005), indicating frequent activation of the DNA damage response in the absence of BLM. It has also been shown that BS cells display elevated levels of loss of heterozygosity (LOH), due to exchanges between homologous chromosomes (LaRocque et al., 2011; Luo et al., 2000; Suzuki et al., 2016).

Although SCEs can be used as a surrogate marker for collapsed forks and DSBs, their locations could until recently only be mapped cytogenetically at megabase resolution (Aguilera and Gomez-Gonzalez, 2008), which did not allow for investigations of the location and potential causes of fork stalling and collapse in BS. We recently described a single-cell sequencing based technique, Strand-seq, which can be used to map SCEs at kilobase resolution, enabling novel studies of their locations and potential causes (Falconer et al., 2012; Sanders et al., 2017).

Besides its ability to regress replication forks and dissolve Holliday junctions, BLM has been shown to bind and unwind guanine-quadruplex (G-quadruplex, or G4) structures *in vitro* (Sun et al., 1998; Wu et al., 2015). G4 structures are stable secondary DNA structures that form at specific guanine-rich DNA motifs containing 4 stretches of 3 or more consecutive guanine residues, separated by variable length spacers (Bochman et al., 2012; Rhodes and Lipps, 2015). The canonical G4 motifs contains 1-7 nucleotide spacers (G_3+_N_1-7_G_3+_N_1-7_G_3+_N_1-7_G_3+_), but DNA motifs containing shorter G-stretches or longer spacers can also fold into G4 structures. G4s are known to constitute barriers for replication fork progression (Lopes et al., 2011) and we hypothesized that BLM deficiency leads to replication fork stalling, collapse, and SCE formation at sites of G4s. Here we show that SCEs in BLM deficient cells indeed occur frequently at sites of G4 motifs, especially when G4 motifs are present in transcribed genes. Our results suggest that the elevated SCE rates in BS cells are mainly caused by failure to resolve G4 structures ahead of replication forks, and not by a higher frequency of crossovers resulting from differences in the resolution of Holliday junctions or the repair of DSBs. Furthermore, we show that loss of heterozygosity (LOH) events appear to be more frequent in BLM deficient cells, as was previously reported for murine cells (Luo et al., 2000), although these events were exceedingly rare in our study. We propose that besides LOH, recombination at G-quadruplexes occurring in transcribed genes is a major contributor to genome instability and cancer predisposition in Bloom syndrome patients.

## RESULTS

### EBV-transformed cells display SCE hotspots in common fragile sites independent of BLM status

We previously reported that elevated SCE rates in BS cells are detected by Strand-seq and that SCE rates in either WT or BS cells are not affected by the culture of cells in the presence of the thymidine analogue bromodeoxyuridine (BrdU) (van Wietmarschen and Lansdorp, 2016). BrdU incorporation is required for SCEs detection by cytogenetics as well as Strand-seq. To address the question of whether SCEs occur at random or at specific locations in the genome, we performed Strand-seq on a panel of 8 different cell lines, 4 obtained from healthy donors (two primary fibroblast and two EBV transformed B-lymphocyte cell lines) and 4 cell lines from BS patients (two fibroblast and two B-cell lines) (see Table S1 for details). We generated several hundred single-cell Strand-seq libraries from each cell line and confirmed that BS cells display roughly ~10 times higher SCE levels than their WT counterparts (Figure 1A, 1B, S1A, and S1B). Although the resolution at which we could map individual SCEs varied widely (Figure 1C and S1C), the median SCE mapping resolution was approximately 10Kbp, and >95% of all SCE could be mapped to regions smaller than 100Kb for each cell line (Table S1).

**Figure 1.**
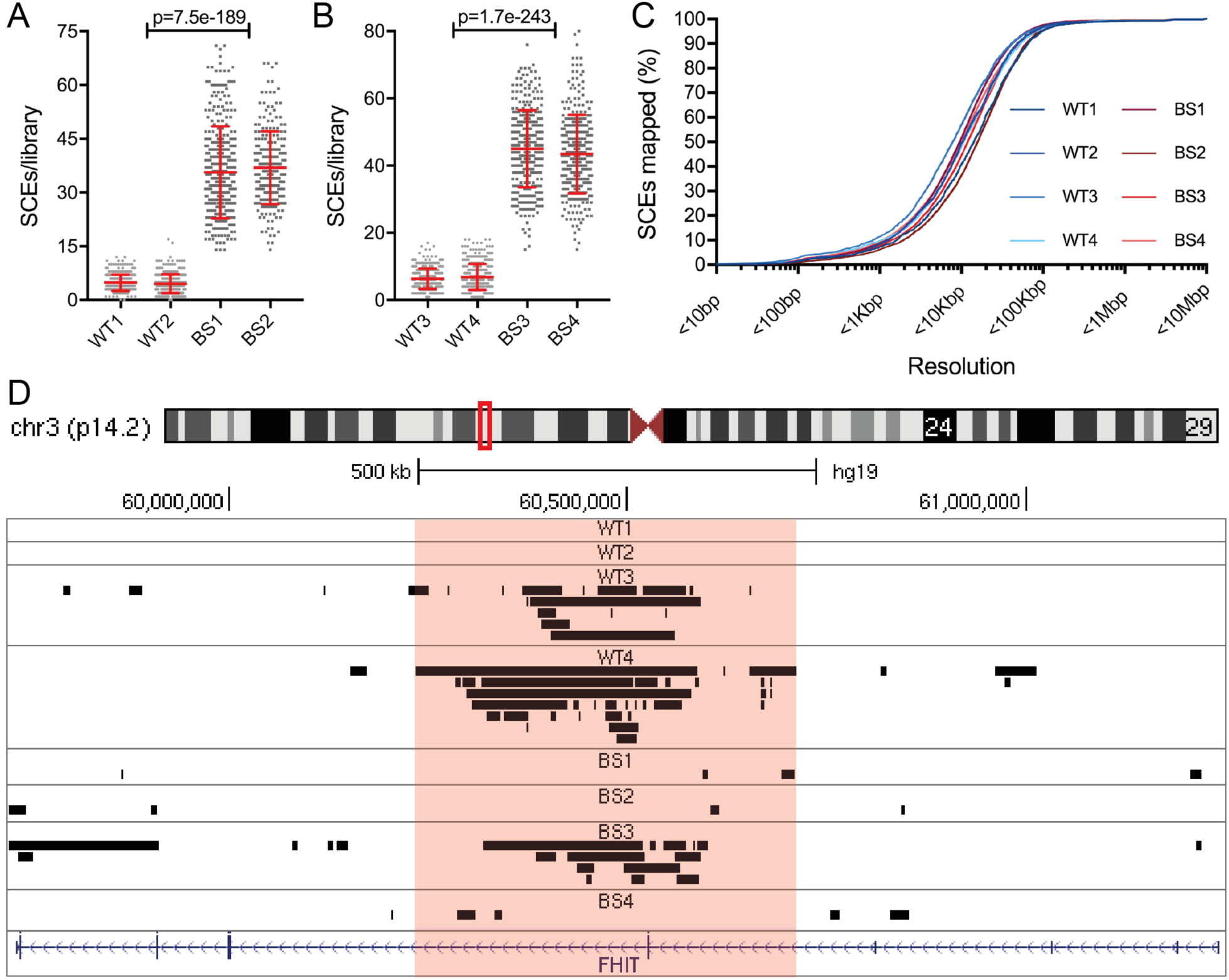
High resolutions SCE mapping reveals hotspots in common fragile sites. (A,B) SCE rates detected in primary fibroblasts (A) and EBV-transformed B-lymphocytes (B) obtained from healthy donor and BS patients. Each grey point represents number of SCEs detected in a single-cell Strand-seq library, red lines indicate mean +/- standard deviation. P-values were calculated using ANOVA. (C) SCE mapping resolutions across all 8 cell lines. Lines represent percentage of the total number of SCEs mapped at resolutions below indicated values. (D) Example of SCE hotspot detected within FHIT gene (FRA3B). Mapped SCE regions for each cell line were uploaded onto the UCSC Genome Browser. Black bars represent genomic locations of SCE regions; size indicates mapping resolution using the BAIT program. Red box indicates the location of SCE hotspot as detected in 3/4 EBV-transformed cell lines.

We next studied the distribution of SCEs across the genome. A strong correlation between chromosome size and the number of SCEs on each chromosome was found in both normal and BS cells (Figure S1D and S1E), as one would expect if SCEs were randomly distributed on a global level. However, several chromosomal regions were found to display higher than expected numbers of overlapping SCE regions, for example within the known fragile site FRA3B in the FHIT gene on chromosome 3p14.2 (Figure 1D). SCE hotspots occurring in known common fragile site (CFS) were detected in all the EBV-transformed B-lymphocyte cell lines, but none of the primary fibroblast lines (Table 1). CFSs are sites of the genome known to display elevated levels of DNA breaks under conditions of replication stress, which can be induced by treatment with the DNA polymerase inhibitor aphidicolin or as a result of viral transformation (Durkin and Glover, 2007). Over a hundred aphidicolin-induced CFSs and many more rare fragile sites occurring in any specific genetic background have been described (Durkin and Glover, 2007). We detected SCE hotspots in a total of 10 CFSs, and in 6 cases the CFS SCE hotspot was only detected in one cell line. Whether this reflects a specific genomic predisposition of the individual from whom the EBV line was derived or a somatic alteration in these cells is not known. Of note, no SCE hotspots were detected in any of the fibroblast cell lines, suggesting that this phenotype is intrinsic to EBV-transformed B-lymphocytes. Although BS cells display a 10-fold higher SCE rate than WT cells, the frequency of SCEs in CFSs is remarkably similar between all EBV-transformed cell lines, with approximately 2-9% of libraries showing an SCE within a given hotspot. This suggests that BLM plays a minor role in the processing of stalled or collapsed replication forks at common fragile sites (Table 1).

**Table 1.** SCE hotspots in common fragile sites. Overview of all SCE hotspots detected in human WT and BS cell lines, as well as frequency of SCE occurrence within hotspots. SCE hotspots were only detected in EBV-transformed cell lines, and all hotspots occurred within known common fragile sites.

| CFS | Coordinates | Genes(s) | Cell line | Libraries with SCE in hotspot | Total Nr of SCEs | P-value |
| --- | --- | --- | --- | --- | --- | --- |
| **FRA1A** | chr1:10,628,012-10,771,196 | PEX14, CASZ1 | BS3 | 13/326 (4.0%) | 14667 | 4.2e-9 |
| **FRA3B** | chr3:60,235,449-60,601,404 | FHIT | WT3 | 15/334 (4.5%) | 2128 | 3.2e-18 |
|  |  |  | WT4 | 28/320 (8.8%) | 2326 | 1.5e-42 |
|  |  |  | BS3 | 13/326 (4.0%) | 14667 | 1.3e-4 |
| **FRA5H** | chr5:58,828,828-59,116,208 | PDE4D | WT4 | 10/330 (4.0%) | 2326 | 3.8e-10 |
| **FRA7B** | chr7:5,880,409-6,120,310 | CCZ1, PMS2 | WT3 | 11/334 (3.3%) | 2128 | 4.8e-13 |
|  |  |  | BS4 | 11/306 (3.6%) | 12724 | 5.0e-5 |
| **FRA7B** | chr7:6,707,575-6,967,206 | PMS2CL, CCZ1B | WT3 | 29/334 (8.7%) | 2128 | 8.9e-50 |
|  |  |  | BS4 | 19/306 (6.2%) | 12724 | 2.9e-14 |
| **FRA7J** | chr7:69,775,932-69,923,390 | AUTS2 | WT4 | 8/330 (2.4%) | 2326 | 8.2e-9 |
| **FRA10G** | chr10:52,320,213-52,640,424 | ASAH2B, A1CF | BS3 | 14/326 (4.3%) | 14667 | 3.2e-6 |
| **FRA11G** | chr11:103,881,556-104,600,858 | PDGFD | BS4 | 17/306 (5.6%) | 12724 | 2.4e-5 |
| **FRA16D** | chr16:78,463,763-78,719,484 | WWOX | WT4 | 8/330 (2.4%) | 2326 | 3.6e-7 |
|  |  |  | BS3 | 13/326 (4.0%) | 14667 | 2.8e-6 |
| **FRA17q21** | chr17:43,517,064-43,840,133 | PLEKHM1, LRRC37 | BS4 | 10/306 (3.3%) | 12724 | 6.7e-3 |

### Bloom syndrome SCEs are enriched in transcribed genes

We next investigated the distribution of SCEs relative to specific genomic features of interest (FOIs). While the resolution of SCE mapping using Strand-seq is orders of magnitude better compared to cytogenetics, the method is not able to map them at base-pair resolution. In over to overcome this limitation, we developed a custom algorithm that compares the distribution of observed SCEs locations in relation to a genomic feature of interest (FOI) to simulated random distributions relative to such a FOI (see Methods section for details). For each cell line, we performed a permutation analysis to calculate the frequency of actual SCE regions overlapping with a given FOI and compared it against the expected background frequency. This analysis yields relative SCE enrichments with a FOI and allows for statistical assessment of the strength of the association.

We first turned to transcribed genes in view of the proposed mechanisms by which transcriptional activity can lead to genome instability and mutations (Gaillard and Aguilera, 2016). Firstly, collisions between the transcription and replication machineries can lead to replication fork stalling, collapse, and DNA breakage (Gaillard and Aguilera, 2016; Helmrich et al., 2013). Secondly, co-transcriptional R-loops could also form a potential barrier to the replication fork and cause fork stalling and collapse (Aguilera and Gaillard, 2014; Sollier and Cimprich, 2015). BLM unwinds R-loops and the absence of BLM has been linked to genome instability at sites of R-loops (Grierson et al., 2013). To study a possible link between SCE locations and transcriptional status we studied the transcriptional activity in each of our 8 cell lines using RNA-seq. Genes were divided into two categories based on the number of fragments per kilobase of processed transcript per million fragments mapped (FPKM) values: active (transcribed) genes (FPKM>1) and silent (non-transcribed) (FPKM<1) genes, resulting in an average of 60% (~23.000) of all genes classified as active and 40% (~16.000) as silent.

A significant enrichment of SCE regions overlapping with gene bodies was found in all BS cells, but in none of the WT cell lines (Figure 2A). However, these enrichments were apparently not affected by gene activity, as both active and silent genes showed the same relative SCEs enrichment (Figure S2A and S2B). To ensure that the higher number of SCEs in the BS cell lines did not affect our analysis, we subsampled the SCE regions from each cell line to match the cell line with the lowest number of SCEs (WT1). In this case, we again detected significant SCE enrichments for overlap with genes in all the BS cell lines, but not in any of the WT cell lines (Figure S2C), indicating that transcription by itself does not appear to play a strong role in SCE formation.

**Figure 2.**
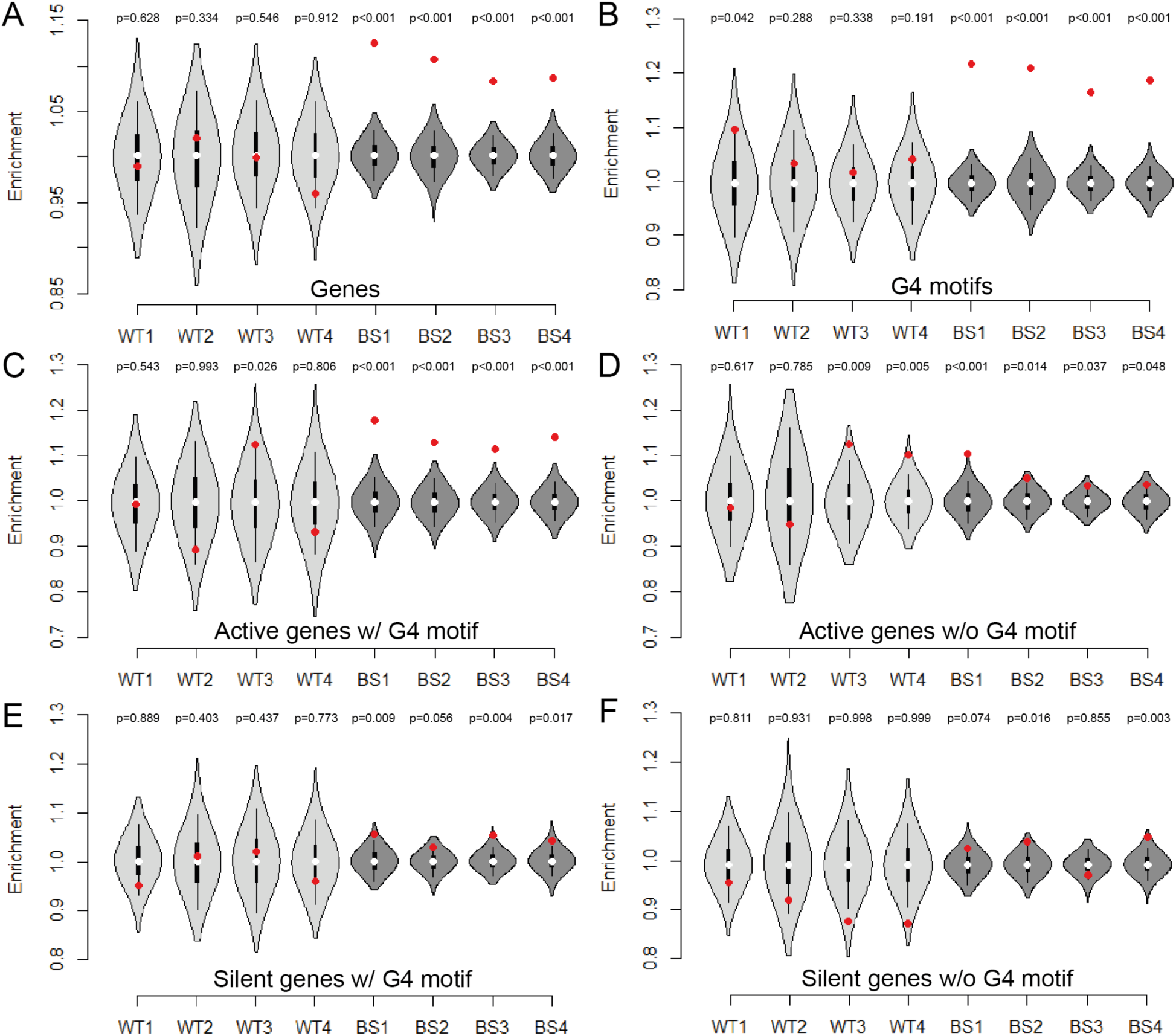
Bloom syndrome SCEs are enriched at G4 motifs and in active genes containing G4 motif(s). Relative SCE enrichments (red points) over random distributions (violin plots) for SCEs overlapping one or more genes (A), G4 motif G_3+_N_1-7_G_3+_N_1-7_G_3+_N_1-7_G_3+_ (B), active genes containing one or more G4 motifs (C), active genes without G4 motifs (D), silent genes containing one or more G4 motifs (E), and silent genes without G4 motifs (F). All values were normalized to the median permuted value and relative SCE enrichments over these values were plotted on the y-axis. P-values indicate the proportion of permuted overlaps equal to or higher than overlap with observed SCE regions.

### Bloom syndrome SCEs are enriched at sites of G-quadruplex motifs

We next considered the possibility that the intragenic SCE enrichments might be partially caused by the presence of G-quadruplexes in or around genes. BLM is known to bind and unwind G4 structures *in vitro* (Huber et al., 2006; Sun et al., 1998; Wu et al., 2015), and G4 motifs occur frequently within gene bodies and promoters (Eddy and Maizels, 2008; Huppert and Balasubramanian, 2007). To assess SCE enrichments at G4 motifs, we determined the distributions of both the canonical and alternative G4 motifs over the genome using a custom algorithm, and performed our SCE enrichment analysis on these regions. As expected, BS SCE regions displayed significant enrichments at G4 motifs, while WT SCE regions did not (Figure 2B).

Because single-stranded DNA (ssDNA) arising during transcription is believed to promote the formation of G4 structures (Bochman et al., 2012), we hypothesized that presence of G4 motifs in genes was the cause of the observed SCE enrichments in BS cells. We therefore divided all genes into 4 categories based on 1) transcriptional activity and 2) the presence or absence of intragenic G4 motifs, and performed a separate SCE enrichment analysis for each category. Active gene bodies containing at least one G4 motif showed the strongest enrichment in BS cells (Figure 2C); weaker enrichments were observed in active genes without G4 motifs (Figure 2D) and silent genes with G4 motifs (Figure 2E) and SCEs were not significantly enriched in silent genes lacking G4 motifs (Figure 2F). SCEs were also significantly enriched in the gene promoter region of BS cells (Figure S2D). However, this enrichment did not appear to depend on the transcriptional status of the associated genes (Figure S2E and S2F). Taken together our results point to a synergistic effect of transcriptional activity and the presence of G4 motifs in genes on the enrichment of SCEs in BS cells.

For our SCE enrichment analysis at G4 motifs, we used a stringent 10Kb size cutoff for SCE regions because G4 motifs occur frequently (at ~8.6Kb intervals on average) and including larger SCE regions would result in increased noise in our analysis because of the high likelihood of (permuted) SCE regions overlapping G4 motifs purely due to their size. Using less stringent size cutoffs did indeed decrease the relative SCE enrichment values for the BS cells, but not the WT cells (Figure S3A), although BS SCE enrichments remained significant for all cutoffs used. To further confirm that SCE enrichments at G4 motifs are specific for the BS cell lines, we added increasingly large flanking sites to SCE regions in order to increase random overlap with G4 motifs, and presumably decrease SCE enrichments. We did indeed detect an inverse relationship between SCE enrichments and size of the flanking regions, but only in the BS cell lines (Figure S3B). This suggests that BS SCEs do indeed frequently occur at G4 motifs, while WT SCEs are randomly spread over the genome with respect to positions of G4 motifs.

We also detected BS SCE enrichments at sites containing G4 motifs containing smaller (n1-3) and larger (n1-12) spacer regions (Figure S3C, S3D) although in both cases the enrichments are weaker than for the canonical G4 motif. This suggests that either the canonical G4 motif is more likely to form G4 structures *in vivo*, or that BLM displays some level of specificity for unwinding G4 structures with medium-sized loops. Significant SCE enrichments were also detected at previously described “observed quadruplex regions” (Figure S3E) reported to constitute all regions in the genome capable of forming quadruplex-like structures (Chambers et al., 2015). As before, SCE enrichments were not affected by SCE subsampling (Figure S3F).

To further ensure that SCEs enrichment was specific to G4 motifs, we analyzed the relationship between SCEs enrichments and other genomic features such as stretches of high A/T or G/C content. We could exclude that enrichments were caused by nucleotide slippage or high GC content, as SCEs were specifically depleted in genomic regions with A-rich motifs (A_3+_N_1-7_A_3+_N_1-7_A_3+_N_1-7_A_3+_) or high GC content across all eight cell lines (Figure S3G and S3H).

### Bloom syndrome SCE enrichments at putative quadruplexes are strongest on template DNA strands within transcribed genes

Intragenic G4 motifs can occur either on the template (transcribed) or the coding (non-transcribed) strand (Figure 3A) and it is believed that this G4 motif ‘strandedness’ affects how G4 structures influence gene expression (Maizels, 2012). In order to assess if G4 strandedness also affects SCE formation, we separated all G4 motifs into different categories based on whether they occur on template or coding strands in active vs silent genes, and performed SCE enrichment analysis for these locations. Our results indicate that G4 motif strandedness can also affect SCE formation in the absence of BLM. We find that overall, template strand G4 motifs show a higher BS-specific SCE enrichment than coding strand G4 motifs and that this effect is strongest for G4 motifs within active genes (Figure 3B-3E). Strikingly, no SCE enrichments were detected for either template or coding strand G4 motif in silent genes (Figure 3F and 3G), or for intergenic G4 motifs (Figure 3H). These results confirm the synergistic effect of transcriptional activity and the presence of a G4 motif for the formation of a SCE in BS cells. Perhaps the transcribed template strand is more susceptible to the formation of a G4 structure upon transcription which, if unresolved by the BLM helicase, can give rise to fork collapse when the replication machinery encounters this G4 structure. Our results also suggest that R-loops are not the main cause of SCEs in BS cells, as R-loops are believed to be stabilized specifically by G4 motifs occurring on the coding (displaced) strand RNA synthesis (Aguilera and Gaillard, 2014).

**Figure 3.**
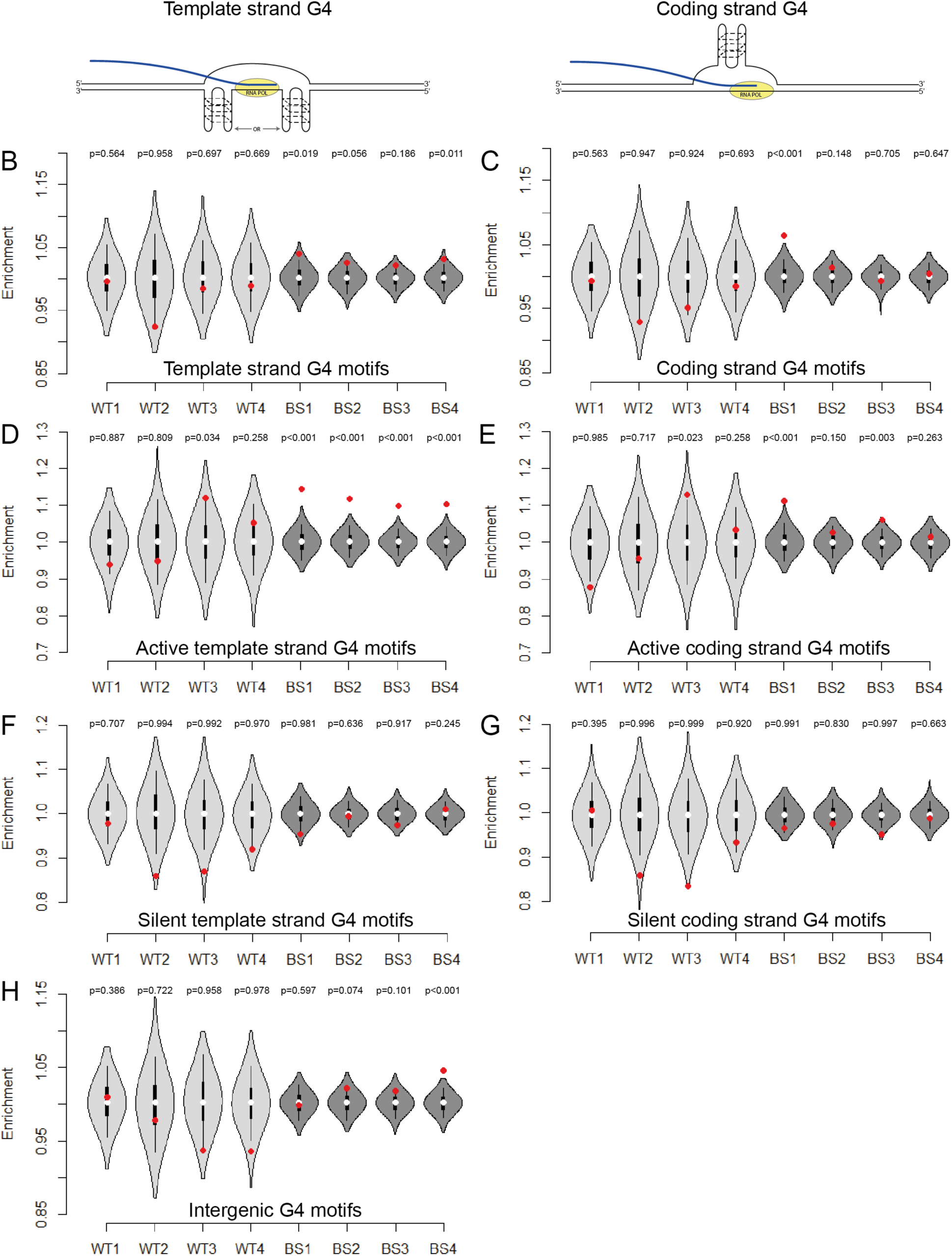
Bloom Syndrome SCE enrichments occur at G4 motifs in transcribed genes. (A) Intragenic G4 motifs can occur on either the template strand (transcribed by RNA polymerase), or on the coding (non-transcribed strand). Template strand G4 structures can theoretically occur ahead of behind RNA polymerase. (B-H) Relative SCE enrichments (red points) over random distributions (violin plots) for SCEs overlapping G4 motifs occurring on intragenic template strands (B), coding strands (C), template strands of active genes (D), coding strands for active genes (E), template strands for silent genes (F), coding strands for silent genes (G), and for intergenic G4 motifs (H). P-values indicate the number of permuted overlaps equal to or higher than overlap with SCE regions.

### Bloom syndrome SCE enrichments at G4 motifs confirmed in Blm^-/-^ mouse embryonic stem cells

To confirm that the BS SCE enrichment patterns we detected are a direct result of BLM deficiency, we next generated *Blm* knockout cells in an F1 hybrid mouse embryonic stem (ES) cell line (129Sv-Cast/EiJ) by means of the Crispr/Cas9 technology. We used different combinations of two guide RNAs to generate loss-of-function mutants by deleting *Blm* exon 19. This exon encodes for part of the HRDC domain, which is critical for BLM’s role in both Holliday junction resolutions (Kim and Choi, 2010) and G4 unwinding (Chatterjee et al., 2014). We selected 3 homozygous and 1 heterozygous clones with the desired deletions and characterized these deletions by Sanger sequencing (Table S2), measured *Blm* mRNA expression levels by qRT-PCR (Figure 4A), and confirmed the elevated SCE rates by Strand-seq (Figure 4B, S4A, and S4B). Interestingly, we detected intermediately high SCE rates in the *Blm*^+/-^ cells, even though previous studies reported that cells from heterozygous family members of BS patients display normal SCE levels (Chaganti et al., 1974; Ellis et al., 1995). SCEs in libraries made from the ES cells could also be mapped at kilobase resolution (Table S2).

**Figure 4.**
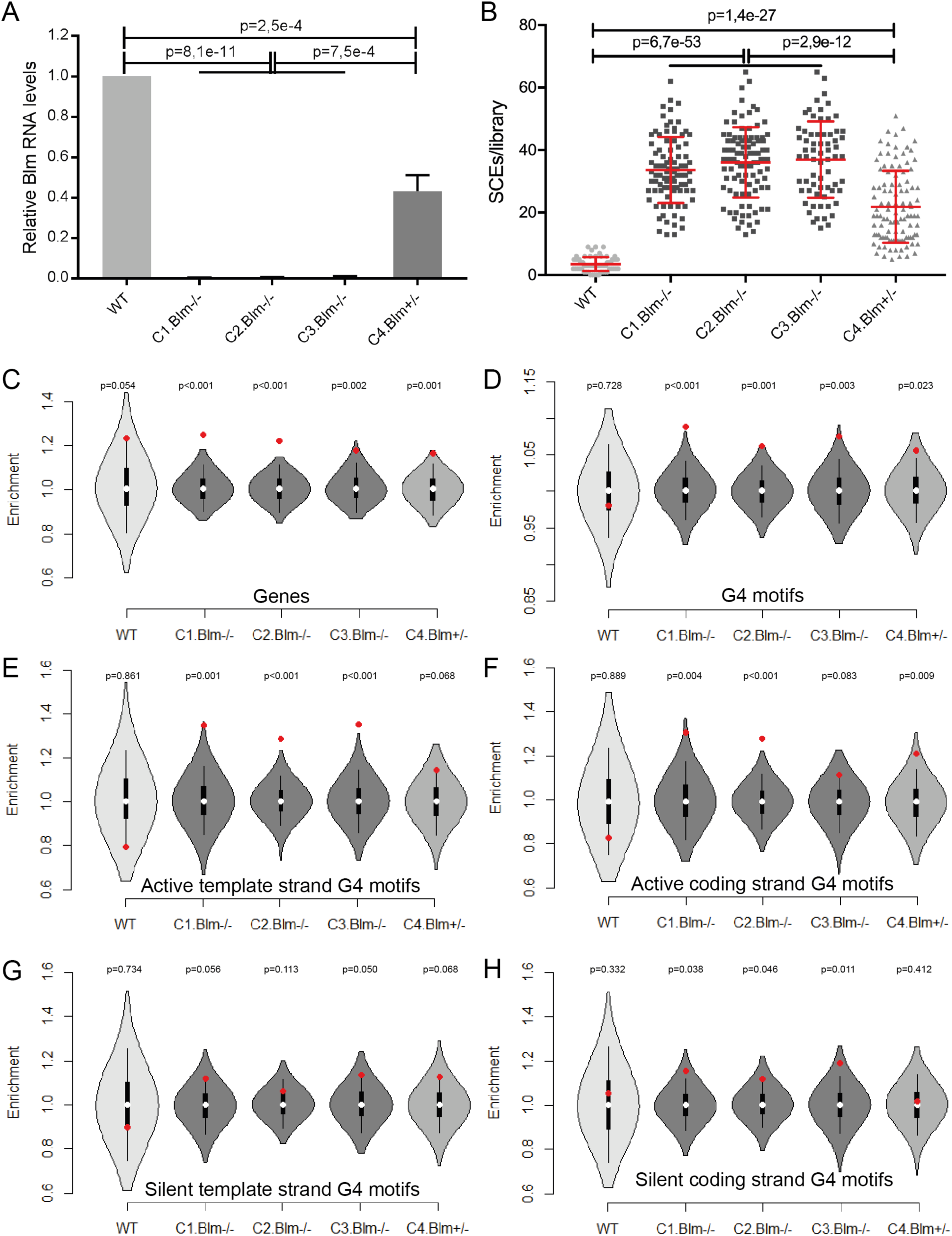
*Blm*^-/-^ mouse ES cells display similar elevated SCE rates and SCE enrichments at G4 motifs to human Bloom syndrome cells. (A) qRT-PCR for Blm transcripts in WT and *Blm* mutant cell lines. Expression levels were normalized to levels in WT cells for each of three replicate experiments. P-values were calculated using t-test and ANOVA. (B) SCE rates detected WT, *Blm*^-/-^, and *Blm*^+/-^ mouse ES cells. Each grey point represents number of SCEs detected in a single-cell Strand-seq library, red lines indicate mean +/- standard deviation. P-values were calculated using t-test and ANOVA. (C-H) Relative SCE enrichments (red points) over random distributions (violin plots) for SCEs overlapping one or more genes (C), G4 motifs (D), and G4 motifs occurring on template strands of active genes (E), coding strands for active genes (F), template strands for silent genes (G), coding strands for silent genes (H). P-values indicate the number of permuted overlaps equal to or higher than overlap with observed SCE regions.

Using the identified SCE regions, we performed the same analysis as described above for the human cell lines. As before, we also generated RNA-seq data for each of ES cell clones to assess the effect of transcriptional activity and G4 strandedness on SCE enrichments. Although we did not detect any clear increased SCE enrichments in genes for the *Blm* mutant cell lines (Figure 4C) or an effect of transcriptional activity (Figure S4C, S4D), we did confirm that these cells display SCE enrichments at canonical G4 motifs (Figure 4D) as well as alternative G4 motifs (Figure S4E, S4F). We detected significant SCE enrichments at sites of G4 motifs occurring on both template and coding strands in the absence of Blm (Figure S4G-H) and confirmed that SCE enrichments in absence of Blm are strongest at G4 motifs occurring on template strands in active genes (Figure 4E), weaker for coding strand G4 motifs in active genes (Figure 4F), and absent in both template and coding strand G4 motifs in silent genes (Figure 4G-H). Like in the human cell lines, we found no SCE enrichments at sites of intergenic G4 motifs (Figure S4I). The F1 hybrid ES cells we used to generate our *Blm* mutants contain >21 million known heterozygous positions, including 72,660 canonical G4 motifs that only occur on one homolog (36,547 in the 129Sv background, and 36,203 in the Cast/EiJ background). To find further evidence of a direct link between G4s and SCEs, we identified all observed SCE regions that overlap a single discordant G4 motif, and identified the homologs that these SCEs occurred on. We found that on average, 69% of informative SCEs in the *Blm*^-/-^ cell lines occurred on the same homolog as the G4 motifs, which is significantly different (p<0.01) from the expected 50% if there was no causal relationship between G4 motifs and SCEs (Figure S4J). No significant deviation from the expected 50/50 ratio was detected in the WT or the *Blm*^+/-^ cell lines. Combined, these results confirm that SCEs mainly form at putative G4 structures in absence of Blm, and especially at those G4s present in the template strands of active genes.

### Blm^-/-^ cells display elevated loss of heterozygosity at very low frequencies

Since SCEs are exchanges of genetic material between identical sister chromatids, they normally do not result in any mutations. However, if an exchange event occurs between homologs instead of sister chromatids, this can lead to loss of heterozygosity (LOH) (Moynahan and Jasin, 1997). LOH is a known driver of cancer, and it has previously been shown that BLM deficient cells display elevated LOH levels (LaRocque et al., 2011; Luo et al., 2000; Suzuki et al., 2016).

We used the high frequency of known single-nucleotide polymorphisms in the F1 ES cells to distinguish homologous chromosomes and identify regions that underwent LOH. To track LOH levels over time, we maintained continuous cell cultures of both the WT and the three *Blm*^-/-^ ESC clones over the course of 30 passages (~75 cell divisions), sorted single cells at different time points (passages 0, 20, and 30), and performed single-cell whole-genome sequencing (scWGS). We then assigned sequencing reads to either genetic background and identified LOH regions as regions displaying loss of reads from one allele coinciding with gain of reads from the other allele. At the same time, we also identified chromosomal and local copy number variations (CNVs) to confirm that LOH regions are not confirmed by deletions, and to determine if the *Blm*^-/-^ cells display aberrant levels of CNVs.

We did not detect a single LOH region in the WT cells at any of the three time points, and only a total of four unique LOH regions in the three *Blm*^-/-^ clones (Figure 5A). Two of these four regions were detected in a single library at a single time point, while the two others were detected at multiple time points and their frequency increased over time. However, the chromosomes harboring these more frequent LOH regions, chr 8 in C1 and chr 1 in C2, were trisomic (Figure S5). Trisomies for chromosomes 1 and 8 were detected in all 4 cell lines, and in each case their frequency increased over time (Figure S5). This suggests that trisomy led to clonal expansion within the cell populations, and the detected LOH regions had no effect on cellular proliferation.

**Figure 5.**
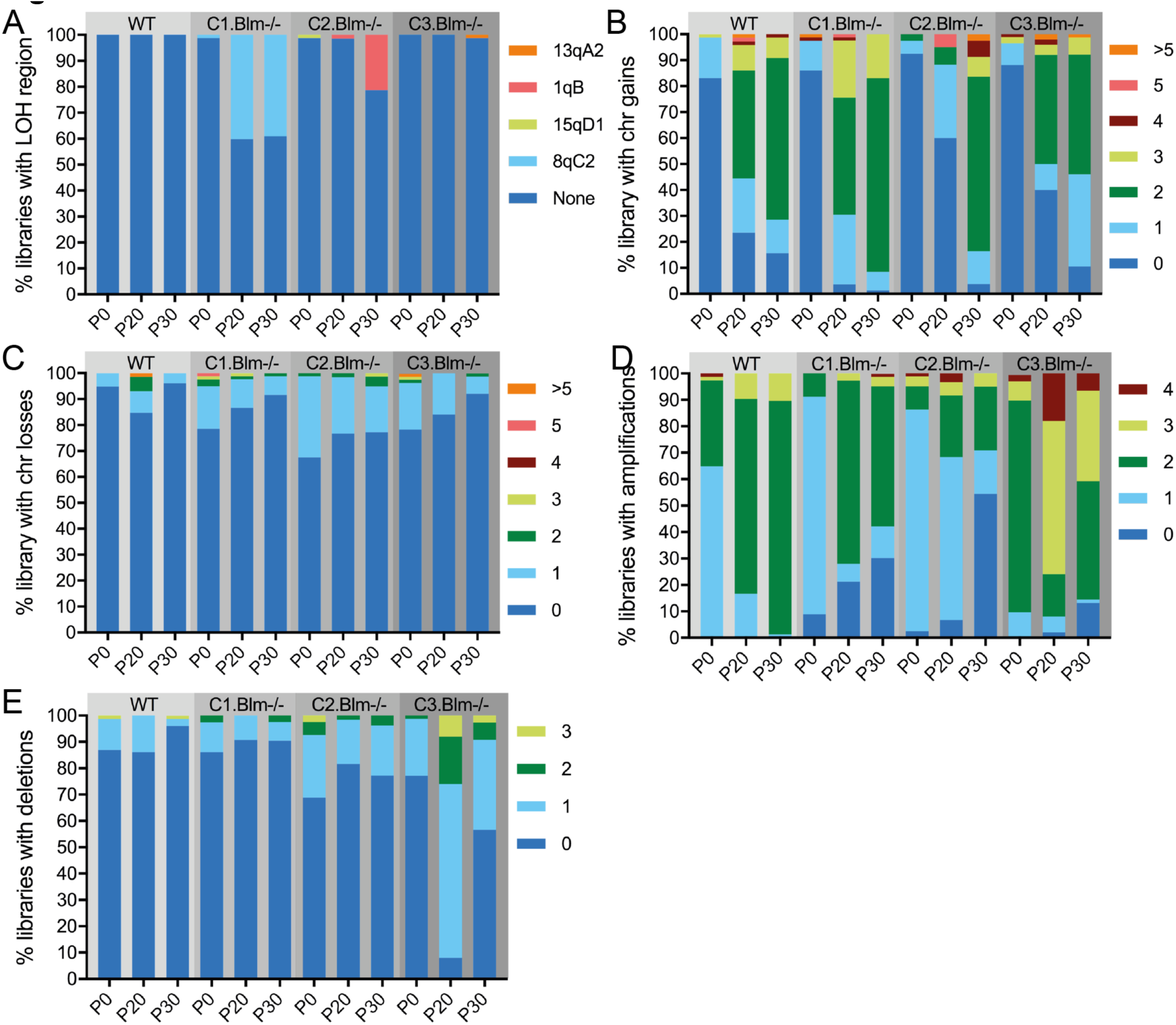
*Blm*^-/-^ mouse ES cells display elevated loss of heterozygosity, no evidence for effect on aneuploidy and local copy number variations. Frequency of unique LOH regions (A), chromosome gains (B), chromosome losses (C), amplifications (D), and deletions (E) detected at in single-cell whole genome sequencing libraries at different passages in WT, *Blm*^-/-^, and *Blm*^+/-^ mouse ES cells.

Overall, the CNV analysis revealed that all four cell lines displayed increasing number of chromosome gains over time, with no clear difference between the WT and *Blm*^-/-^ lines (Figure 5B). We detected relatively low levels of whole chromosome losses and local amplifications and deletions, which did not increase in frequency over time or showed differences between the WT and *Blm*^-/-^ lines (Figure 5C-5E). Taken together, these results indicate that Blm deficiency does not lead to marked elevation of local or large chromosomal CNVs in murine embryonic stem cells.

## DISCUSSION

Elevated SCE rates are a hallmark feature of BS cells (Chaganti et al., 1974; van Wietmarschen and Lansdorp, 2016), but the exact mechanism behind this phenotype is not fully understood. A major obstacle to unravelling potential causes of SCEs was that they could so far only be mapped using standard cytogenetic detection methods (Aguilera and Gomez-Gonzalez, 2008). For this study, we used Strand-seq to map SCEs in both normal and BS cells at kilobase resolution, and we show that SCEs frequently occur at sites of putative G4 structures in both BLM deficient human and murine cells. Consistent with the high frequency of G4 motifs in gene bodies and promoters (Eddy and Maizels, 2008; Huppert and Balasubramanian, 2007), we found strong BS SCE enrichments for both features, indicating that these genomic loci are mutation prone in BS cells. While there does not appear to be a direct effect of transcriptional activity on SCE enrichment patterns, we did detect strong SCE enrichments in BS cells in transcribed genes containing one or more G4 motifs, especially when the G4 motifs occur on intragenic template strands. This is in line with evidence that G4 structures preferentially form in regions of single-stranded DNA (Bochman et al., 2012) and that G4 structures form mainly in euchromatic regions of the genome (Hansel-Hertsch et al., 2016). The fact that we frequently detect SCEs occurring on the same homolog as homolog-specific G4 motifs in our *Blm* mutant ES cells provides further evidence that G4 structures are a direct cause of SCEs in BLM deficient cells.

Our results are consistent with proposed models of BLM unwinding G4 structures during DNA replication (Chatterjee et al., 2014; Croteau et al., 2014) and with previous reports that BLM binds and unwinds G4 structures *in vitro* (Sun et al., 1998; Wu et al., 2015), and facilitates telomere replication by unwinding G4 structures ahead of the replication fork (Drosopoulos et al., 2015). Our results show that BLM is not only required to unwind G4 structures in telomeres, but throughout the genome. Indeed, G4 structures are known to pose barriers for DNA replication (Lopes et al., 2011) and previous studies have shown that specialized helicases such as Dog-1, Pif1 and FANCJ are required to prevent instability at G-rich genomic DNA in *C.elegans* (Cheung et al., 2002), yeast (Paeschke et al., 2011), and man (Castillo Bosch et al., 2014). The fact that such other helicases cannot compensate for loss of BLM suggests that these helicases do not have redundant functions, but are either specific for subsets of G4 structures, or that they cooperate to unwind G4 structures. Such a cooperation has previously been proposed for BLM and FANCJ (Suhasini et al., 2011).

Compared to expected values based on random SCE distributions, we detected significant (P<0.001) ~20% enrichments for SCEs at G4 motifs in the absence of BLM. This is almost certainly an underestimation of the actual SCE enrichments at G4 structures for two reasons. Firstly, a subset of mapped SCE regions will overlap with the relatively abundant G4 motifs purely because of their size, leading to non-specific overlaps and elevated noise levels in the permutation analysis. Using stringent cutoffs for the SCE region size increases relatively enrichment values, but our analysis is still limited by the kilobase resolution of the Strand-seq method to map SCE. Secondly, it has been estimated that cells harbor ~10.000 actual G4 structures (Hansel-Hertsch et al., 2016), compared to ~350.000 canonical G4 motifs predicted *in silico*. Because our analysis is based on SCEs overlapping with G4 motifs, we likely overestimate overlaps with G4 structures in our permutation analysis, leading to reduced enrichment estimates. Taking this into consideration, we believe that the 20% enrichments we detected strongly underestimate actual SCE enrichments at G4 structures, which might be much higher than suggested here.

Although SCEs are typically considered non-mutagenic themselves, they are considered markers for genome fragility and for sites of increased rates of mutations (Bradley et al., 1979). BS SCEs frequently occur in transcribed genes, indicating these sites are subjected higher mutation rates. This is arguably highly detrimental, and elevated intragenic mutations rates are likely to contribute to the strong cancer predisposition associated with Bloom syndrome. This also helps explain a unique feature of Bloom syndrome, which predisposes patients to a wide range of cancers instead of towards specific types of tumors (German, 1993).

It has previously been reported that SCEs frequently occur in CFSs, particularly in cells undergoing replication stress (Glover and Stein, 1987). We confirm the existence of SCE hotspots in CFSs in EBV-transformed B-cells, but not primary fibroblasts. Strikingly, SCE frequencies in CFS hotspots are similar between the WT and BS cell lines instead of being elevated in BS cells along with the global SCE rates. These results contradict established models of BLM actively counteracting SCE formation at stalled replication forks (Wu, 2007), but seem consistent with our hypothesis that BLM specifically prevents replication fork stalling and collapse at G4 structures.

DNA repair via HR can also result in LOH when the homologous chromosome is used for repair instead of the sister chromatid (Moynahan and Jasin, 1997). It has previously been shown that BLM deficient cells display high levels of LOH (LaRocque et al., 2011; Luo et al., 2000; Suzuki et al., 2016). However, these results were obtained using systems that rely on selection of cells that underwent LOH at a specific locus. We could detect LOH regions throughout the genome without selective pressure. We detected 4 unique LOH regions across 3 independent *Blm* mutant clones, each of which was kept in continuous culture for ~75 cell divisions. This number of divisions would result in 3,8x10^22^ offspring cells for each parental cell, compared to an estimated 1,2x10^10^ cells in an adult mouse body. We confirm that *Blm*^-/-^ cells display elevated LOH levels (no LOH regions were detected in the WT cell line), but show that LOH in the *Blm*^-/-^ cells is a rare event that did not seem to provide a proliferative advantage in our experimental setup. Although BLM has been linked to chromosome segregation (Chan et al., 2007) and BLM deficient cells display a higher frequency of micronuclei (Yankiwski et al., 2000), we did not detect higher levels of aneuploidy in the *Blm*^-/-^ cells, indicating that BLM deficiency does not necessarily result in increased levels of aneuploidy.

In conclusion, we provide evidence that the BLM helicase is required for unwinding of G4 structures occurring at sites of transcription to prevent stalling of replication forks at these known barriers for DNA replication. Our results help explain the elevated SCE rates occurring in BS cells and suggest that mutation rates are highest at sites of G-quadruplexes in transcribed genes in BS cells, potentially contributing to the high frequency of cancer seen in patients with Bloom syndrome. As such, we propose that BLM deficiency leads to excessive recombination and chromosomal fragility at sites of G-quadruplexes in transcribed genes. Combined with elevated LOH levels, this chromosome fragility is likely to play a role in the strong cancer predisposition.

## AUTHOR CONTRIBUTIONS

Conceptualization, N.v.W. and P.M.L.; Methodology, N.v.W, N.H., D.C.J.S.; Software and Formal Analysis: N.v.W., V.G.; Investigation: N.v.W., S.M., N.H.; Writing – Original Draft, N.v.W., Writing – Reviewing & Editing, N.v.W., S.M., D.C.J.S., V.G., P.M.L.; Supervision & Funding Acquisition: P.M.L.

## ACKNOWLEDGEMENTS

We thank Dirk Hockemeyer, Marcel van Vugt, and Peter Stirling for critical reading of this manuscript, Inge Kazemier and Karina Hoekstra-Wakker for technical assistance, and Ester Falconer, Mark Hills, and all group members for discussions and feedback. Financial support was provided by an Advanced Grant from the European Research Council to P.M.L.

## FIGURE LEGENDS

**Figure S1.**
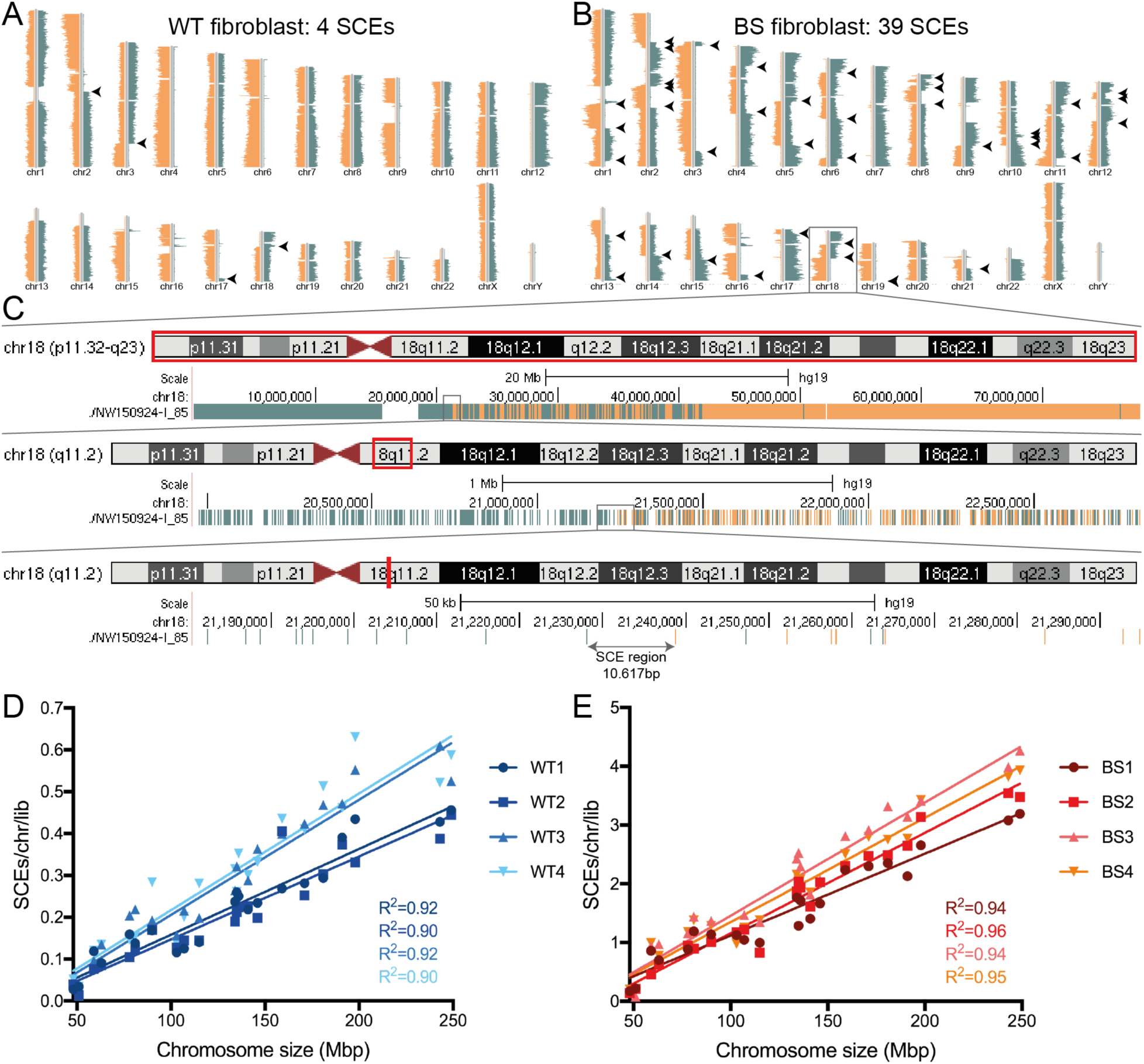
High resolution SCE mapping using Strand-seq. (A,B) Representative Strand-seq libraries generated from a WT fibroblast (A) and a BS fibroblast (B). Mapped DNA template strand reads are plotted on directional chromosome ideograms; Crick (positive, sense) reads in green, Watson (negative, antisense) reads in orange. SCEs are identified as a switch in template strand state, indicated by black arrowheads. (C) Graphical representation of high resolution mapping for SCE in chromosome 18 from Strand-seq library shown in (B). Strand-seq library is uploaded into USCS genome browser, SCE is seen as a switch from two Crick template strands (solid green) to one Watson and one Crick strand (mixed green and orange). Zooming in on approximate SCE region allows for identification of individual reads flanking the switch from Crick to Watson. The SCE is mapped to the region between these reads, in this case a 10.617bp region. The BAIT software package (Hills et al., 2013) was used for automated SCE mapping. This software calculates ratios of Watson/Crick reads in a bin (1 for 100% Watson reads, −1 for 100% Crick reads, and 0 for 50% Watson/Crick), identifies SCEs as changes in this ratio along chromosomal segments, and then localizes SCEs to the locations of the ratio switch based on the first 20 reads flanking each site of the SCE region. (D,E) Correlations between average numbers of SCEs/chromosome/library and chromosome size for WT (D) and BS (E) cells. R^2^ are color-matched to the cell lines.

**Figure S2.**
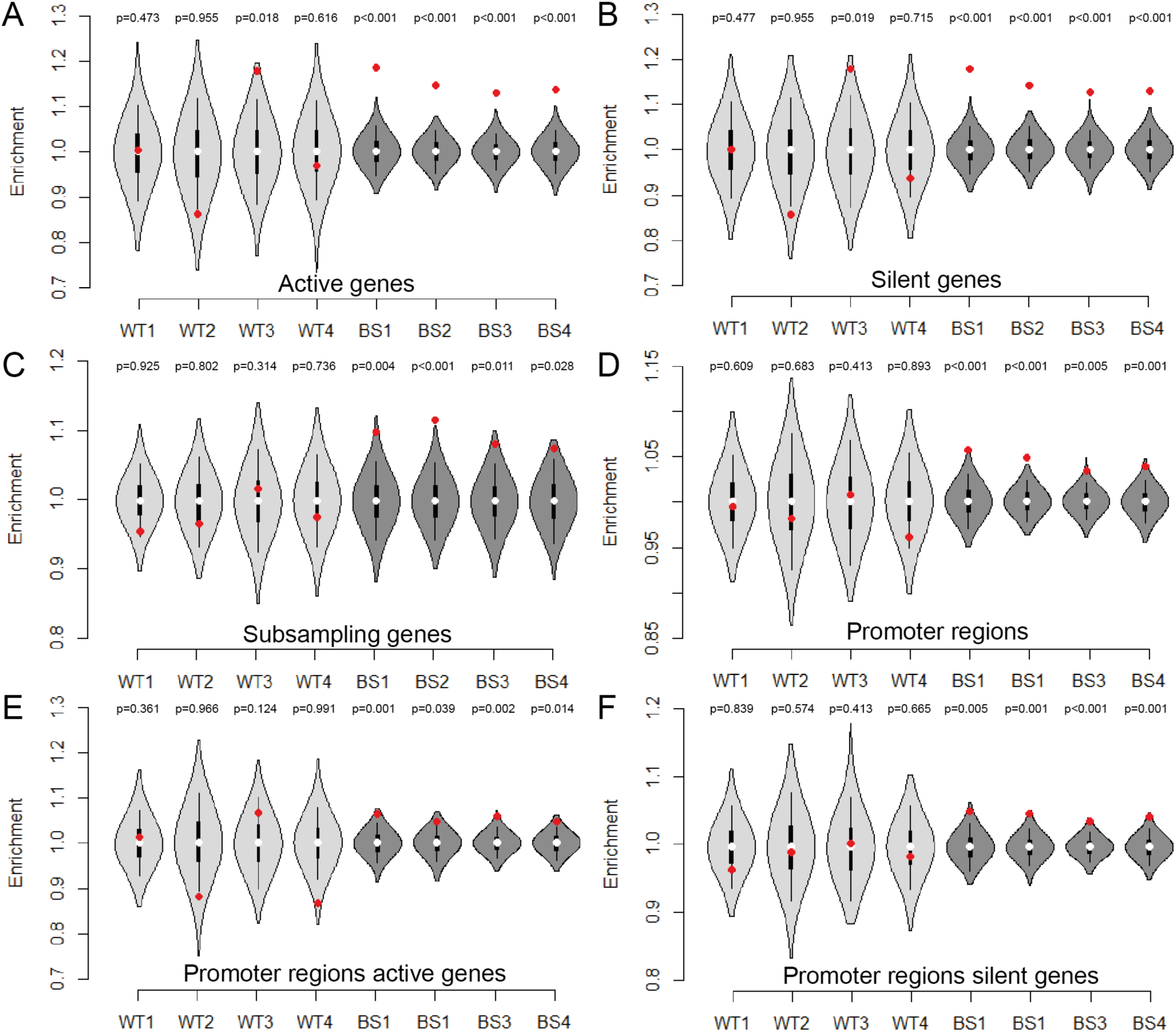
SCE enrichments in genes and promoters, and effect of transcriptional activity. Relative SCE enrichments (red points) over random distributions (violin plots) for SCEs overlapping transcribed genes (A), non-transcribed genes (B), genes after SCE subsampling (C), promoter regions (D), promoter regions associated with transcribed genes (E), and promoter regions associated with non-transcribed genes (F). P-values indicate the number of permuted overlaps equal to or higher than overlap with SCE regions. SCE subsampling was achieved by randomly selecting 1559 SCEs (the number of SCEs mapped in the smallest dataset) for each randomized permutation, as well as for the overlap of experimentally determined SCE regions with genes or promoter regions.

**Figure S3.**
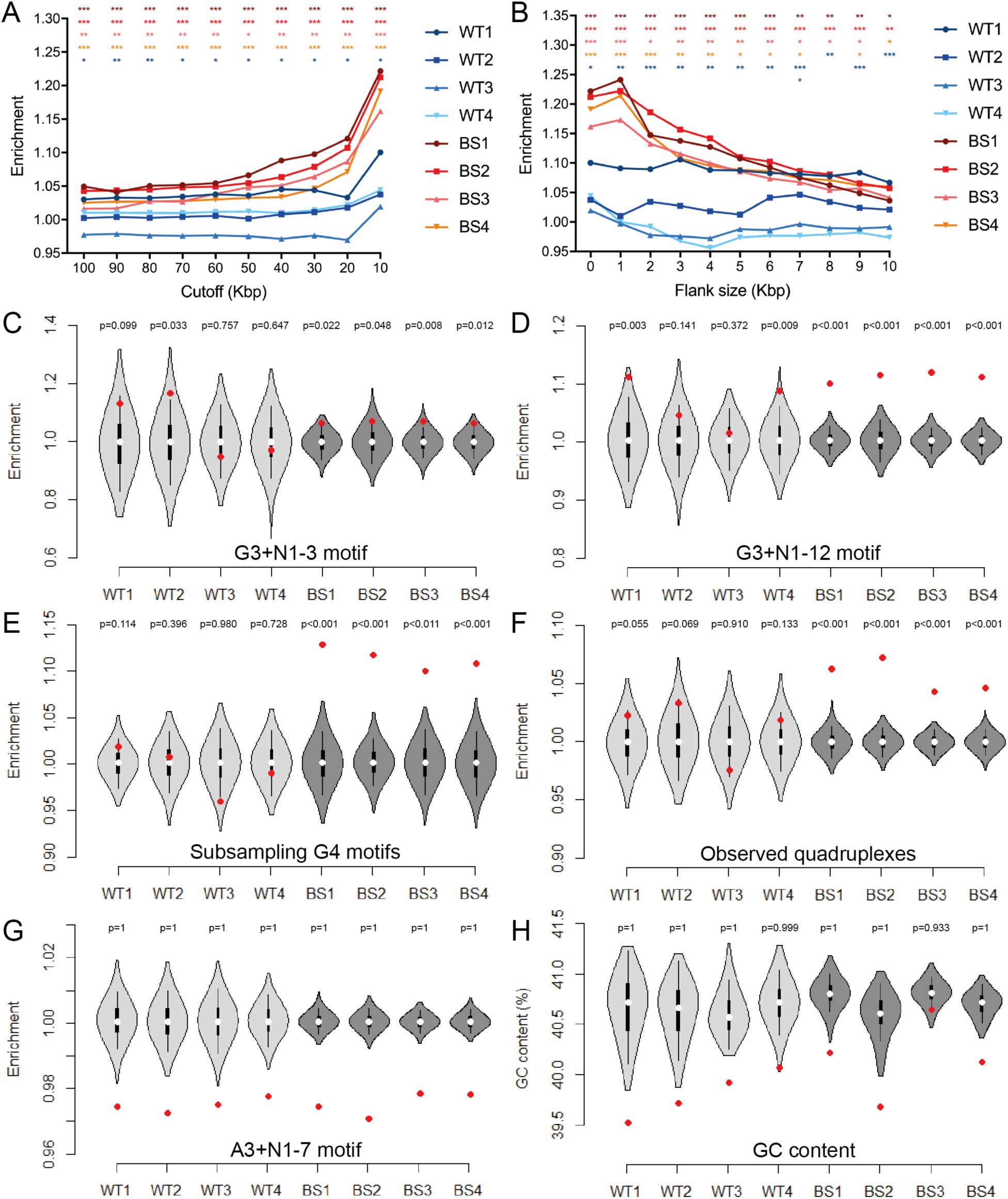
SCE enrichments at alternate G4 motifs, effect of different SCE size cutoffs. (A) Effect of differently size cutoffs for SCE regions included in analysis of relative SCE enrichment over median permuted overlaps. (B) Effect of adding increasingly large flanking regions to <10Kb SCE regions on relative SCE enrichment over median permuted overlaps. *: p<0.05, **: p<0.01, *** p<0.001 for color-matched cell line. (C-F) Relative SCE enrichments (red points) over random distributions (violin plots) for SCEs overlapping the alternate G4 motifs G_3+_N_1-3_G_3+_N_1-3_G_3+_N_1-3_G_3+_ (C), and G_3+_N_1-_12G_3+_N_1-12_G_3+_N_1-12_G_3+_ (D), canonical G4 motifs after subsampling SCE regions (E), and previously reported observed quadruplex forming regions (F). (G) SCE enrichments for A_3+_N_1-7_A_3+_N_1-7_A_3+_N_1-7_A_3+_motif. (H) GC content of all SCE regions per cell line compared to GC content of permuted SCE regions. For enrichment analyses, P-values indicate the number of permuted overlaps equal to or higher than overlap with SCE regions. SCE subsampling was achieved by randomly selecting 1559 SCEs (the number of SCEs mapped in the smallest dataset) for each randomized permutation, as well as for the overlap of experimentally determined SCE regions with genes or promoter regions. An SCE region size cutoff of 10Kb was used for these analyses.

**Figure S4.**
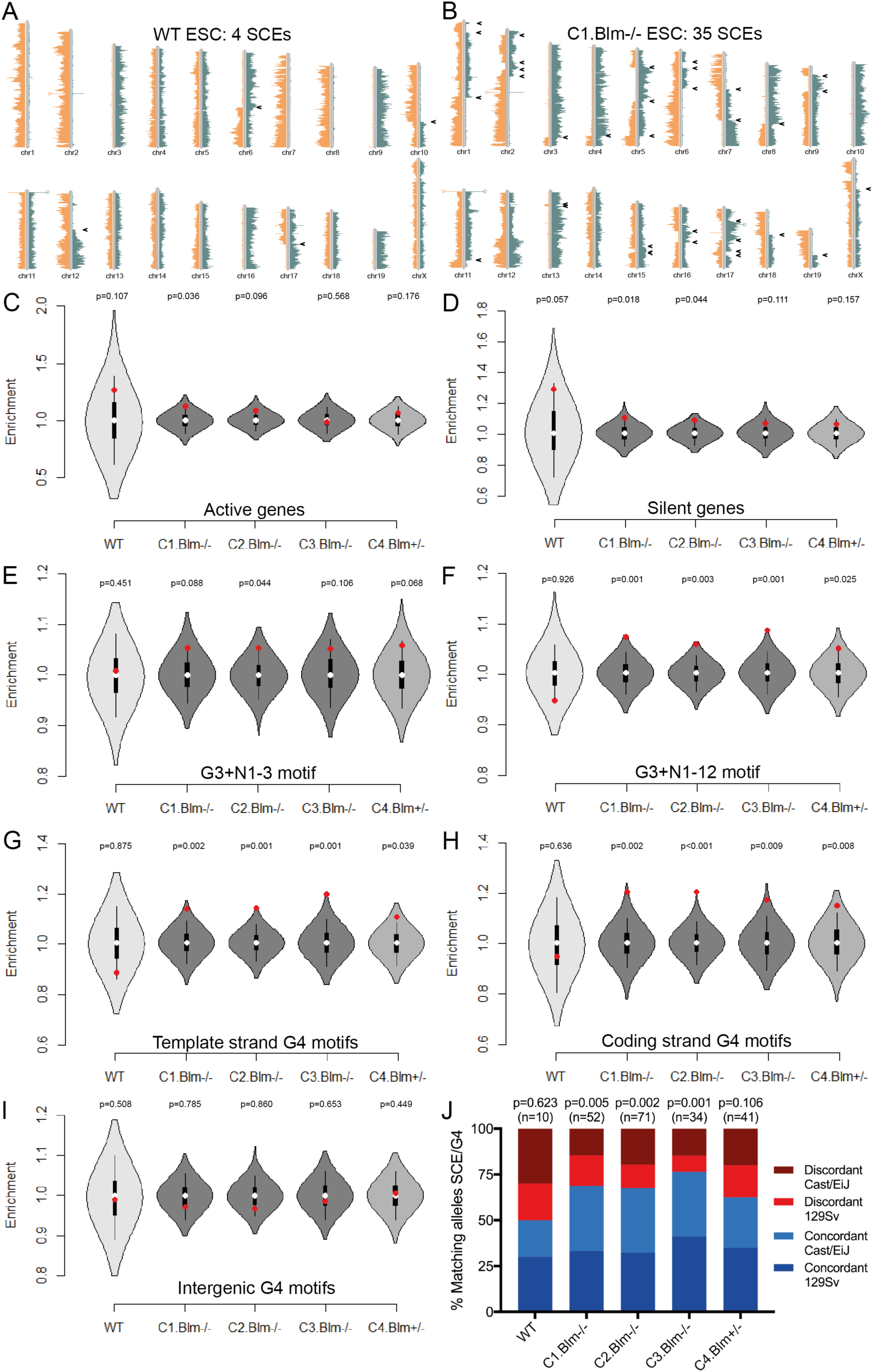
Effect of *Blm* knockout on SCE enrichment at alternate G4 motifs. (A, B) Representative Strand-seq libraries generated from a WT (A) and a C1.*Blm*^-/-^ (B) cell. Mapped DNA template strand reads are plotted on directional chromosome ideograms; Crick (positive, sense) reads in green, Watson (negative, antisense) reads in orange. SCEs are identified as a switch in template strand state, indicated by black arrowheads. (C-I) Relative SCE enrichments (red points) over random distributions (violin plots) for SCEs overlapping on active genes (C), silent genes (D), the alternate G4 motifs G_3+_N_1-3_G_3+_N_1-3_G_3+_N_1-3_G_3+_ (E), and G_3+_N_1-12_G_3+_N_1-12_G_3+_N_1-12_G_3+_ (F), or canonical G4 motifs occurring in intragenic template strands (G), intragenic coding strands (H), or in intergenic regions (I). For enrichment analyses, p-values indicate the number of permuted overlaps equal to or higher than overlap with SCE regions. (J) Frequency of observed SCE regions occurred on the same homolog as allele-specific G4 motifs. Indicated is the homolog containing G4 motif, concordant indicates SCE occurred on same homolog, discordant indicates SCE occurred on opposite homolog. P-values were calculated using binomial distributions based on a 50% chance of SCE and G4 motif occurring on the same homolog.

**Figure S5.**
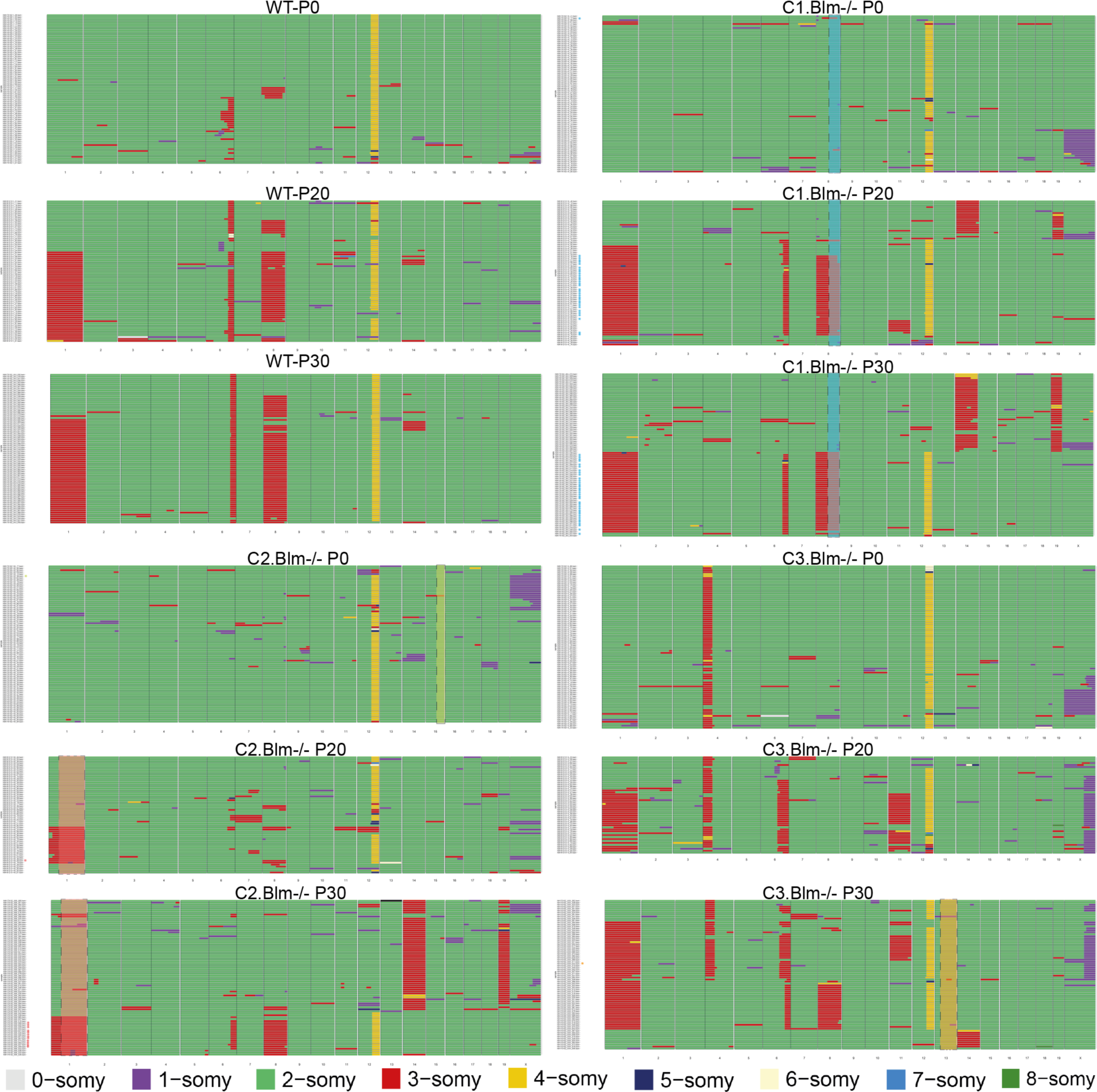
Aneuploidy and CNV analysis in WT and *Blm*^-/-^ ES cell lines. Graphical representation of chromosome copy number states in single-cell whole genome sequencing libraries prepared from WT and *Blm*^-/-^ cells at indicated passage numbers. Each horizontal line represents a single-cell library, with chromosome numbers indicated below each plot. Different colors represent copy number status as detected in 2Mb bins. Cells are clustered based on the similarity of their copy number profile. LOH regions for each population are highlighted (in the same color as Figure 5A), with libraries harboring this LOH region indicated with an asterisk.

**Table S1.** Overview of human cell lines used, Strand-seq libraries made, and SCEs identified and mapped.

| Cell line | Corriell ID | Cell type | Disease state | Strand-seq libraries | Nr of SCEs | Median mapping resolution | SCE regions <100KBp |
| --- | --- | --- | --- | --- | --- | --- | --- |
| **WT1** | GM07492 | Primary fibroblast | Healthy | 321 | 1559 | 15714 | 95,25 |
| **WT2** | GM07545 | Primary fibroblast | Healthy | 354 | 1616 | 11430 | 96,93 |
| **WT3** | GM12891 | EBV-transformed B-lymphocyte | Healthy | 334 | 2128 | 7847 | 96,71 |
| **WT4** | GM12892 | EBV-transformed B-lymphocyte | Healthy | 340 | 2326 | 10722 | 96,28 |
| **BS1** | GM02085 | Primary fibroblast | Bloom Syndrome | 328 | 11156 | 10102 | 97,58 |
| **BS2** | GM03402 | Primary fibroblast | Bloom Syndrome | 229 | 8453 | 17497 | 96,05 |
| **BS3** | GM16375 | EBV-transformed B-lymphocyte | Bloom Syndrome | 326 | 14667 | 13064 | 96,94 |
| **BS4** | GM17361 | EBV-transformed B-lymphocyte | Bloom Syndrome | 306 | 12724 | 10805 | 95,99 |

**Table S2.** Overview of Blm mutant mouse ES cell lines generated and used for this study.

| Clone | Allele | Deletion coordinates | Deletion size | Exon 19 status | Strand-seq libraries | Nr of SCEs | Median mapping resolution | SCE regions <100KBp |
| --- | --- | --- | --- | --- | --- | --- | --- | --- |
| **WT** | Cast | Not mutated | n.a. | WT | 98 | 244 | 8.048bp | 99.16% |
|  | 129 | Not mutated | n.a. | WT |  |  |  |  |
| **C1.Blm<sup>-/-</sup>** | Cast | 71163-71636 | 473bp | Lost | 91 | 3048 | 7.392bp | 98.59% |
|  | 129 | 71167-71396 | 229bp | Lost |  |  |  |  |
| **C2.Blm<sup>-/-</sup>** | Cast | 71166-72195 | 1029bp | Lost | 97 | 3801 | 6.251bp | 98.01% |
|  | 129 | 71166-72195 | 1029bp | Lost |  |  |  |  |
| **C3.Blm<sup>-/-</sup>** | Cast | 71161-71570 | 409bp | Lost | 67 | 2597 | 6.722bp | 98.51% |
|  | 129 | 71161-71570 | 409bp | Lost |  |  |  |  |
| **C4.Blm<sup>+/-</sup>** | Cast | 71377-71848 | 471bp | Lost | 107 | 2437 | 7.554bp | 97.58% |
|  | 129 | Not mutated | n.a. | WT |  |  |  |  |

## METHODS

### Cell cultures

The following cell lines were obtained from the Corriell Cell Repository: GM07492 and GM07545 (primary fibroblasts, normal), GM02085 and GM03402 (primary fibroblasts, Bloom Syndrome), GM12891 and GM12892 (EBV-transformed B-lymphocytes, normal), GM16375 and GM17361 (EBV-transformed B-lymphocytes, Bloom Syndrome). The WT hybrid mouse ES cell line F121.6 (129Sv-Cast/EiJ) was a kind gift from Joost Gribnau (Erasmus University, Rotterdam, the Netherlands). Fibroblasts were cultured in DMEM (Life Technologies) supplemented with 10% v/v FBS (Sigma Aldrich) and 1% v/v penicillin-streptomycin (Life Technologies), B-lymphocytes in RPMI1640 (Life Technologies) supplemented with 15% v/v FBS and 1% v/v penicillin-streptomycin.

ES cells were cultured on mitotically arrested mouse embryonic fibroblast cells in DMEM (Life Technologies), supplemented with 15% v/v FBS (Bodinco BV), 1% v/v penicillin-streptomycin, 1% v/v non-essential amino acids (Life Technologies), 50µM 2-mercaptoethanol (ThermoFisher Scientific), and 1000U/ml leukemia inhibitor factor (Merck). All cells were cultured at 37°C in 5% CO_2_. For Strand-seq, BrdU (Invitrogen) was added to exponentially growing cell cultures at 40µM final concentration. Timing of BrdU pulse was 12 hours for ES cells, 18 hours for fibroblast cell lines, and 24 hours for B-lymphocyte cell lines.

### Generation of Blm mutant ES cell lines

Blm mutants were generated using CRISPR/Cas9 genome editing. sgRNAs were designed to cleave the *Blm* gene at sites flanking exon 19 (see Key Resources Table) and cloned into PX459 plasmid using previously reported protocol (Ran et al., 2013). Combinations of two plasmids (30µg each) were transfected into F121.6 cells by means of electroporation (Biorad Genepulser XL). Cells were incubated for 24 hours before puromycin (1µg/ml) was added to cell culture medium. After 48 hours of selection, resistant colonies were left to grow, picked and expanded. Screening for *Blm* mutant clones was performed by allele-specific PCR of genomic region containing putative deletion.

### qRT-PCR

Exponentially growing cells were harvested and RNA was isolated using the Nucleospin RNA kit (Macherey Nagel). Reverse transcription was performed using Superscript II Reverse Transcriptase (Invitrogen) with random hexamers (Invitrogen). qPCR was performed using SYBR Green I Master (Roche) on the LightCycler480 (Roche).

### Strand-seq and single-cell WGS library preparation

Protocols for single-cell sorting, Strand-seq library preparation (Sanders et al., 2017), and single-cell WGS library preparation (van den Bos et al., 2016) were previously described. For each experiment, 96 libraries were pooled and 250-450bp sized fragments were isolated and purified. DNA quality and concentrations were assessed using the High Sensitivity dsDNA kit (Agilent) on the Agilent 2100 Bio-Analyzer and on the Qubit 2.0 Fluorometer (Life Technologies).

### RNA-seq library preparation

Exponentially growing cells were harvested and RNA was isolated using the Nucleospin RNA kit (Macherey Nagel). RNA-sequencing libraries were prepared using the NEBNext Ultra RNA Library Prep kit for Illumina (NEB) combined with the NEBNext rRNA Depletion kit (NEB). cDNA quality and concentrations were assessed using the High Sensitivity dsDNA kit (Agilent) on the Agilent 2100 Bio-Analyzer and on the Qubit 2.0 Fluorometer (Life Technologies).

### Illumina sequencing

Clusters were generated on the cBot (HiSeq2500) and single-end 50 bp reads (Strand-seq and RNA-seq) or paired-end 150 bp reads (scWGS) were generated were generated using the HiSeq2500 sequencing platform (Illumina).

### Bioinformatics

#### Genome alignment

Indexed bam files were aligned to human (GRCh37) or mouse genomes (GRCm38) using Bowtie2 (Langmead and Salzberg, 2012) for Strand-seq and scWGS libraries, and STAR aligner (Dobin et al., 2013) for RNA-seq libraries.

#### Sister chromatid exchange detection

SCE were identified and mapped with the BAIT software package (Hills et al., 2013), using standard settings. Because BAIT also detects stable chromosomal rearrangements, events that occurred at the exact same locations in >5% of cells from one cell line were excluded from the analysis. SCEs were assigned to homologs by splitting .bam files into separate files for each genetic background based on reads covering informative polymorphisms, and using BAIT to identify on which homologs SCEs occurred.

#### Detection and analysis of SCE hotspots

BAIT-generated .bed files containing the locations of all mapped SCEs were uploaded to the USCS genome browser and hotspots were identified as regions containing multiple overlapping SCEs. P-values were assigned to putative SCE hotspots using a custom R-script based on capture-recapture statistics. Briefly, the genome was divided into bins of the same size as the putative hotspot, and the chance of findings the observed number of SCEs in one bin was calculated based on the total number of SCEs detected in the cell line.

#### Enrichment analysis

A custom Perl script was used for the permutation model. For each of 1000 permutations we generated a random number *n* and shifted all SCEs downstream by *n* bases on the same chromosome. To prevent small-scale local shifts, we required *n* to be a random number between 2 Mbp and 50 Mbp. If the resulted coordinate exceeded chromosome size we subtracted the size of chromosome, so that the SCE is mapped to beginning part of the chromosome, as if the chromosome was circular. We also excluded all annotated assembly gaps before our analysis, to prevent permuted SCE mapping to one of the gap regions. We then determined the number of SCEs overlapping with a feature of interest in each permutation, as well as the original SCE regions. All values were normalized to the median permutated value in order to determine relative SCE enrichments over expected, randomized distributions and to allow for comparison of the different cell lines. Significance was determined based on how many permutations showed the same or exceeding (enrichment) or the same or receding (depletion) overlap with a given genomic feature compared to overlap between the original SCEs and the same feature. Any experimental overlap that lies outside of the 95% confidence interval found in the permutations has a p-value below 0.05 and was deemed significant. Experimental overlaps lying outside of the permuted range were given a p-value below 0.001 as there was a less than 0.1% (1/1000) chance of such an overlap occurring by chance. Enrichment analyses for G4 motifs were performed using a 10Kb SCE region size cutoff, enrichment analysis for genes and promoter regions used a 100Kb size cutoff, unless specified otherwise. Genome and gene annotations were obtained from Ensembl release 75 (GRCh37 assembly, http://www.ensembl.org). Gene bodies were defined as regions between transcription start sites and transcription end sites, gene promoters as 1Kbp regions upstream of transcription start sites. Putative G-quadruplex motifs were predicted using custom Perl script by matching genome sequence against following patterns: G_3+_N_x_G_3+_N_x_G_3+_N_x_G_3+_, where x could be the ranges of 1-3, 1-7, or 1-12 bp. *RNA-seq analysis:* mapped reads were aligned and quantified using STAR aligner (Dobin et al., 2013). FPKM (Fragments Per Kilobase Of Exon Per Million Fragments Mapped) values were calculated for all genes and based on these genes were assigned active (FPKM >1) or silent (FPKM <1) status.

#### Aneuploidy and CNV detection

Aligned libraries were analyzed as previously described using AneuFinder R package (Bakker et al., 2016) using the following settings: low-quality alignments (MAPQ<10) and duplicate reads were excluded, and read counts in 2Mb variable-width bins were determined with a 10-state Hidden Markov Model (HMM) with copy-number states: zero-inflation, null-, mono-, di-, tri-, tetra-, penta-, hexa-, septa- and octasomy.

#### LOH detection

Reads were aligned to either 129Sv or Cast/EiJ genetic background based on covered SNPs. Reads lacking informative SNPs were discarded. 129Sv reads were assigned a positive (Crick) orientation, Cast/EiJ reads a negative (Watson) orientation. The resulting .bam files were analyzed using BAIT and LOH events were detected as switches from mixed background to pure 129Sv or Cast/EiJ background in the absence of deletions (as detected using AneuFinder).

